# Coordinated changes in a cortical circuit sculpt effects of novelty on neural dynamics

**DOI:** 10.1101/2023.10.21.563440

**Authors:** Shinya Ito, Alex Piet, Corbett Bennett, Séverine Durand, Hannah Belski, Marina Garrett, Shawn R. Olsen, Anton Arkhipov

## Abstract

Recent studies have found dramatic cell-type specific responses to stimulus novelty, highlighting the importance of analyzing the cortical circuitry at the cell-type specific level of granularity to understand brain function. Although initial work classified and characterized activity for each cell type, the specific alterations in cortical circuitry—particularly when multiple novelty effects interact—remain unclear. To address this gap, we employed a large-scale public dataset of electrophysiological recordings in the visual cortex of awake, behaving mice using Neuropixels probes and designed population network models to investigate the observed changes in neural dynamics in response to a combination of distinct forms of novelty. The model parameters were rigorously constrained by publicly available structural datasets, including multi-patch synaptic physiology and electron microscopy data. Our systematic optimization approach identified tens of thousands of model parameter sets that replicate the observed neural activity. Analysis of these solutions revealed generally weaker connections under novel stimuli, as well as a shift in the balance e between SST and VIP populations. Along with this, PV and SST populations experienced overall more excitatory influences compared to excitatory and VIP populations. Our results also highlight the role of VIP neurons in multiple aspects of visual stimulus processing and altering gain and saturation dynamics under novel conditions. In sum, our findings provide a systematic characterization of how the cortical circuit adapts to stimulus novelty by combining multiple rich public datasets.

## Introduction

Detecting novel stimuli and adapting to new situations enable animals to identify potential threats or opportunities and increase their chances of survival. Therefore, it is not surprising that novelty powerfully modifies patterns of neural activity to affect perception (Rust & Cohen, 2022; Schomaker & Meeter, 2012), facilitate learning (Bunzeck & Düzel, 2006; Lisman & Grace, 2005), and elicit appropriate behavioral reactions (Sokolov, 1963). Neurons often decrease their response to repeated stimuli and recover the response when a novel stimulus is presented (Bendixen et al., 2012; Henson & Rugg, 2003; Homann et al., 2022; Knight, 1996; Näätänen et al., 1982; Ringo, 1996; Tapper & Molas, 2020; Weber et al., 2019; Xiang & Brown, 1998). However, recent studies showed that different neuronal subclasses are engaged in distinct and diverse ways by stimulus novelty (Garrett et al., 2020, 2023; Pérez-González et al., 2005). For instance, Garrett et al. (2020) demonstrated that vasoactive intestinal polypeptide (VIP) expressing inhibitory cells in the mouse visual cortex exhibited increased responses when presented with novel versus familiar images, suggesting a flexible mode of processing under different conditions. A follow-up study (Garrett et al., 2023) further detailed how stimulus novelty affects specific cell types and circuits, suggesting that novelty acts as a key driver of cellular functional diversity. Elucidating mechanisms that determine neural cell type specific effects of novelty in reshaping neural circuits is crucial for developing an understanding of how the brain operates under diverse conditions encountered by behaving animals.

In the primary visual cortex (V1), one candidate mechanism for explaining the observed changes in neural dynamics due to stimulus novelty is the alteration in the balance between inhibition and disinhibition of excitatory (Exc) neurons. This proposed motif involves Exc neurons that are inhibited by somatostatin (SST) expressing inhibitory neurons, which have mutual inhibition with VIP inhibitory neurons (Fu et al., 2014; Pfeffer et al., 2013). Under this hypothesis, when VIP neurons are activated by novelty, they inhibit the SST neurons, ultimately disinhibiting the Exc neurons. This idea of disinhibition was explored in a modeling study, which successfully reproduced the novelty effect in V1 neurons by introducing spike-timing-dependent plasticity to modulate the inhibitory-to-excitatory connections (Schulz et al., 2021). Although this motif provides a framework for understanding some aspects of neural adaptation to novelty, it is likely part of a more complex network of interactions that remains to be fully understood.

While the above-mentioned motif provides a valuable framework, it’s important to recognize that the effects of stimulus novelty are far from monolithic. For example, novel experiences that share some commonality with past ones, and those that have bare minimum relationships to the past ones are thought to activate distinct neuromodulatory circuits for memory consolidation (Duszkiewicz et al., 2019). In addition, omission of a stimulus is considered a special type of stimulus novelty and is often studied separately from other forms of novelty (Bendixen et al., 2012; Braga & Schönwiesner, 2022). Although these individual types of novelty have been studied over decades, how they interact with each other and how they affect the cortical circuitry has not been well understood.

Recently, new systematic and extensive datasets have been released that characterize visual responses in the mouse brain to both familiar and novel stimuli (Garrett et al., 2023) (https://portal.brain-map.org/explore/circuits/visual-behavior-2p; https://portal.brain-map.org/explore/circuits/visual-behavior-neuropixels), offering excellent opportunities to study novelty effects. The data were collected using 2-photon calcium imaging or extracellular electrophysiology with Neuropixels probes (Jun et al., 2017; Siegle et al., 2021) in awake, behaving mice. These experiments measured neural activity in response to multiple types of stimulus novelty. These include changes in image identity following the repeated presentation of a constant image; omissions of a stimulus from an expected sequence; and absolute stimulus novelty, characterized by exposure to a previously unencountered stimulus. The data demonstrated that cortical neurons exhibit coordinated dynamics in response to each of these novelty types. In this study, we focus on the changes in neuronal dynamics that are observed when mice are exposed to absolute novelty. Specifically, we observed: (1) an increase of the responses of Exc, parvalbumin (PV), and VIP neurons, and (2) altered temporal dynamics in VIP neurons, characterized by slow ramping activity during the omission of the stimulus in familiar conditions, and reduction of such activity when the stimulus is novel (Fig. 1C). These observations create a unique opportunity to study the interaction of different types of novelty at cell subclass level of granularity. Building upon the foundation of these new observations, we conducted the present study to understand what kind of changes in the underlying circuitry may support the observed substantial modifications of cortical dynamics with novelty.

**Figure 1:**
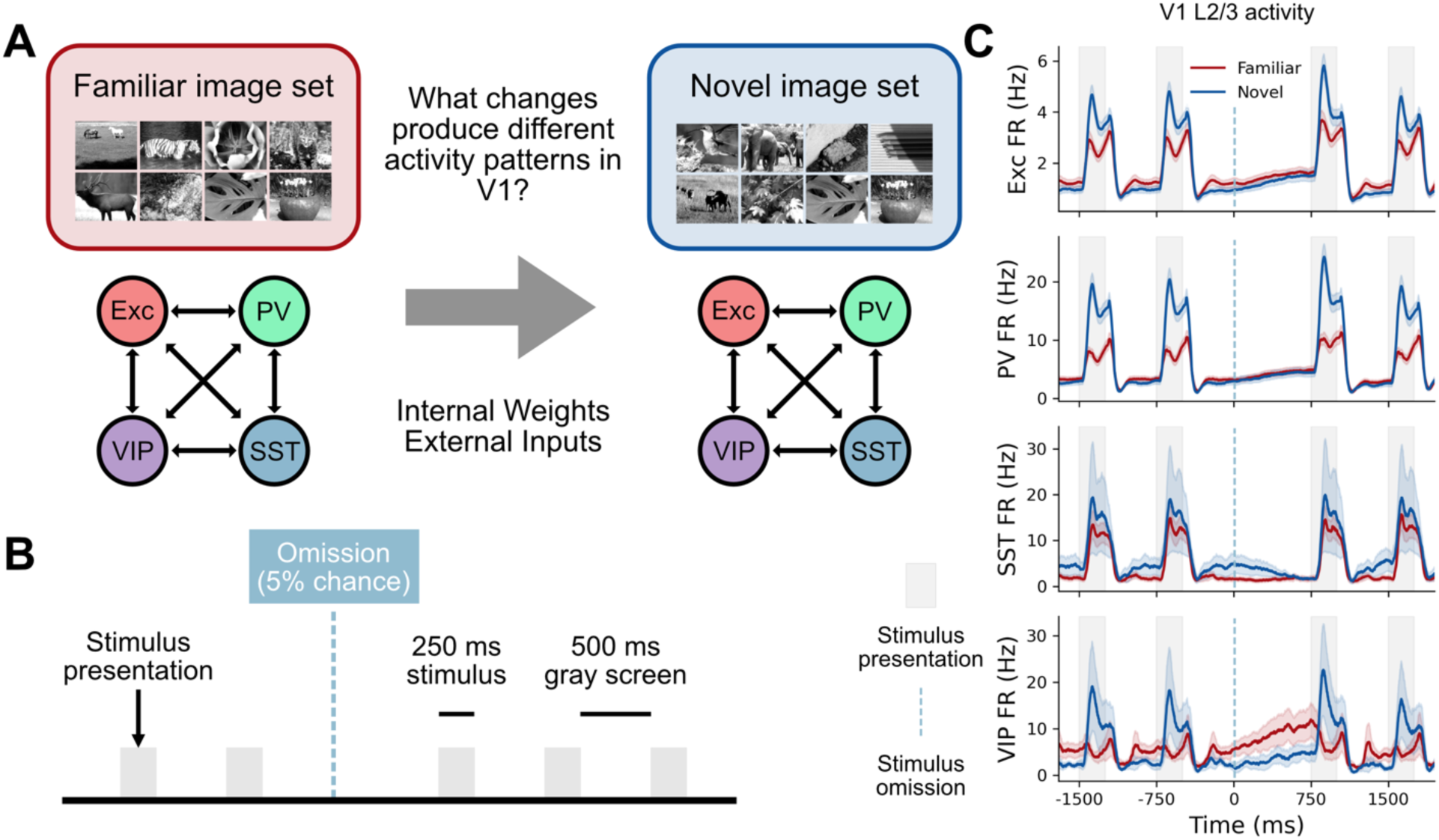
Illustration of the experimental setup and the phenomenon under investigation. A: Schematics of the questions addressed in this study. The image set changes between the Familiar and Novel sessions. B: Essential structure of the task relevant to this study. Stimuli are repeated at a regular cadence (250 ms on, 500 ms off). The stimulus is omitted 5% of the time, but the temporal cadence is preserved. C: Average neural population activity in layer 2/3 around the stimulus omission. The shaded area represents the SEM, with the denominator being the number of specimens. (See Methods for details.)

To achieve this goal, we created network models of neural cell subclass population activity, focusing on the circuit in layer 2/3 (L2/3) of V1. We fit these models to neural activity recorded before and after exposure to absolute novel stimuli and investigated differences in parameters between the networks fit to these Familiar and Novel conditions. We used a stabilized supralinear network (SSN) model (Rubin et al., 2015) for this purpose, which is a population model that can exhibit various nonlinear behaviors (Kraynyukova & Tchumatchenko, 2018; Millman et al., 2020). To enhance the model’s realism, we systematically integrated publicly available datasets describing the architecture of the circuit, including a multi-patch synaptic physiology dataset (Campagnola et al., 2022; Seeman et al., 2018) (https://portal.brain-map.org/explore/connectivity/synaptic-physiology) and electron microscopy dataset (The MICrONS Consortium et al., 2021) (https://www.microns-explorer.org/cortical-mm3), to constrain the model parameters.

Our systematic optimization approach found tens of thousands of sets of parameters that reproduce the observed activity while keeping the parameters within the ranges constrained by structural data. This collection of the parameters was further analyzed to determine the trend of changes in the parameters between the Familiar and Novel conditions. The transition from the Familiar to Novel condition is characterized by general weakening of the connections accompanied by the shift of balance between the SST and VIP populations. Along with it, PV and SST populations experienced overall more excitatory influences compared to Exc and VIP populations. We also identified that the inputs into the VIP population had a unique role in separating image responses and non-image responses and found that the networks in Novel conditions have generally higher gain, but earlier saturation in response to the input stimulus amplitude. In sum, our study offers a quantitative analysis of the V1 L2/3 circuit elements in response to different types of novelty and their interaction, at the cell subclass level of granularity. This work establishes a foundation for understanding the mechanisms that allow the brain to adapt to new information, paving the way for more targeted research.

## Results

### Stimulus novelty influences the responses of V1 neurons to image presentations and omissions

The primary goal of this modeling study was to investigate how mechanistic changes within the network, such as in the connectivity strength, give rise to the complex shifts in the dynamics of the neurons between Familiar and Novel conditions. For this purpose, we utilized the Allen Institute’s visual behavior Neuropixels dataset (https://portal.brain-map.org/explore/circuits/visual-behavior-neuropixels) containing data on neural dynamics in multiple populations of cortical neurons under familiar and novel conditions, which were used to develop our modeling approach as described below.

In the present study, we focused on modeling layer 2/3 of V1 because it is where Garrett et al. (2020, 2023) first observed extensive, cell subclass-specific changes in dynamics in response to absolute novelty. The motivations for using the Neuropixels data instead of the original data from Garrett et al. (2020, 2023), which used two-photon calcium imaging, are the former’s advantage in temporal resolution and direct access to firing rates of individual neurons.

The experiments were conducted with animals trained to respond to changes in image identity. Initial recordings were performed with a set of images used during the training phase (Familiar condition; Fig. 1A). After a recording session with the Familiar image set, the set of images was replaced with a new set to introduce absolute novelty (Novel condition; Fig. 1A). In both Familiar and Novel sessions, the displayed image was occasionally (5% of the time) omitted and replaced with a gray screen (Fig. 1B).

Fig. 1C highlights the differences in the activity patterns of each population between the Familiar and Novel conditions. The major changes include an increase of the firing rates in response to stimulus image presentations (shaded areas) in Exc, PV, and VIP populations (especially, the VIP population shifts from being inhibited by stimuli to responding to stimuli), and reduction of the ramping activity of the VIP population in response to stimulus omission (*t* = 0) in the Novel relative to the Familiar condition (Fig. 1C bottom panel). In the previous publications (Garrett et al., 2020, 2023), the effect of image identity changes has also been discussed. However, we did not include those in our modeling framework because it will require a model to have multiple channels of image inputs and a mechanism that changes the response to the images within the same condition, thus complicating the model. Besides, responses to such contextual novelty have been already described using synaptic plasticity (Aitken et al., 2023; Schulz et al., 2021). Therefore, we selected data segments around omission as the target data for our model, as these observations underscore the complex interplay between the circuit elements in V1, particularly in the interaction between absolute and omission novelty.

### Data-driven SSN model successfully reproduces experimentally observed neural dynamics in Familiar and Novel conditions

To capture the observed neural dynamics, we used a stabilized supralinear network (SSN) model (Rubin et al., 2015) of L2/3 populations. The framework for our model is outlined in Fig. 2. The model consists of four populations: Exc, PV, SST, and VIP. The model’s parameters, most of which are connectivity parameters (influence matrix), are derived from experimental data, including multi-patch synaptic physiology data (Campagnola et al., 2022) and MICrONS electron microscopy data (The MICrONS Consortium et al., 2021). The relative strengths of the connections between populations, along with their errors, are estimated by the following relationship:

**Figure 2:**
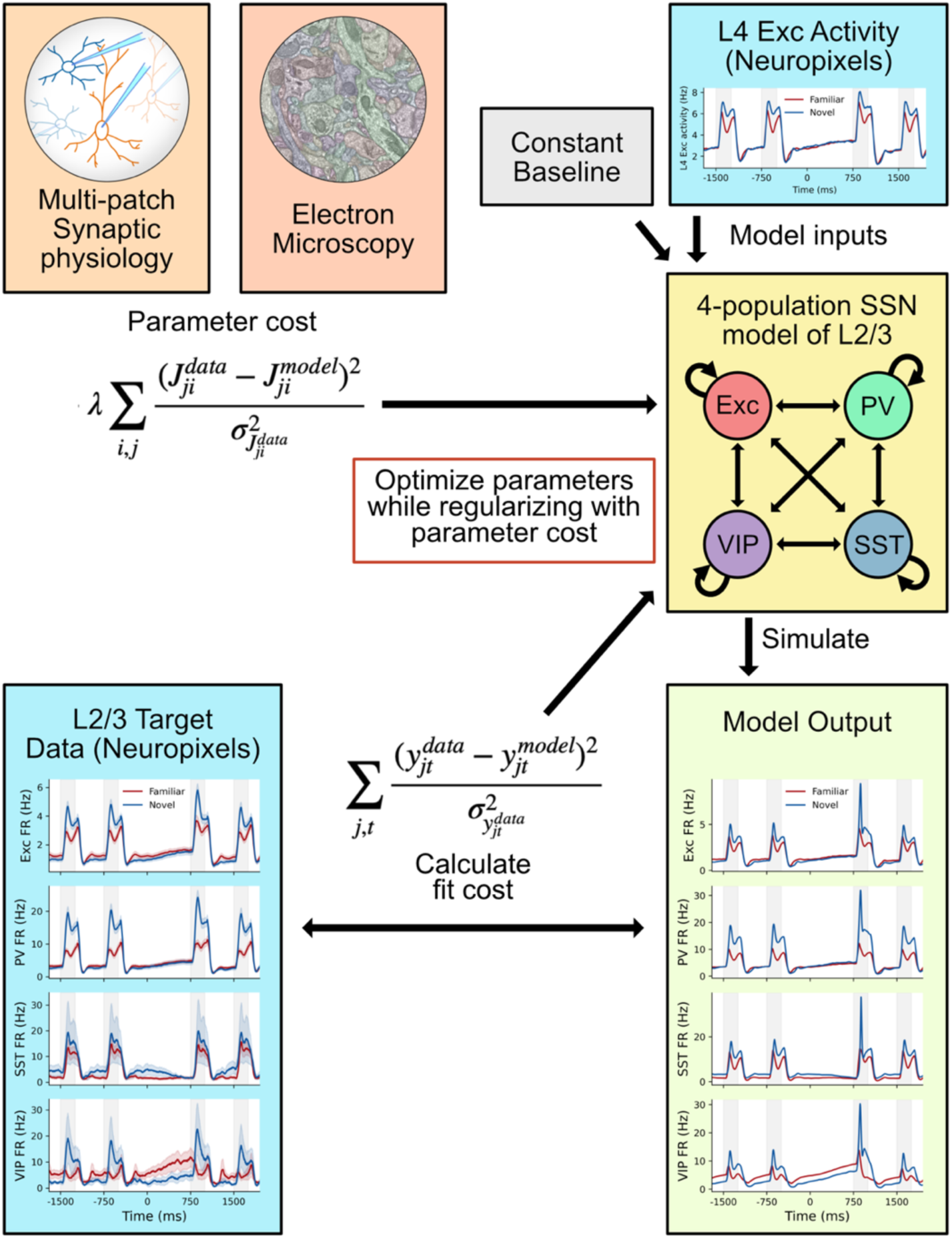
Schematic representation of the modeling workflow. The four-population stabilized supralinear network (SSN) is used to examine the effect of novelty on cortical dynamics. The multi-patch synaptic physiology dataset (Campagnola et al., 2022; Seeman et al., 2018) (https://portal.brain-map.org/explore/connectivity/synaptic-physiology) and MICrONS electron microscopy dataset (The MICrONS Consortium et al., 2021) (https://www.microns-explorer.org/cortical-mm3) are used to construct a parameter cost that constrains the model connectivity parameters. The activity of the L4 Exc population from the Neuropixels data serves as the input drive (together with the baseline constant input), and the model output is compared with L2/3 activity from the Neuropixels to construct the fit-cost. The optimization is repeated 100,000 times with different random initial parameters for each session to gather statistics on the solutions.

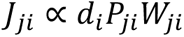

where *J*_*ji*_ is the influence matrix element from population *i* to population *j*, *d*_*i*_ is the density of the neurons of population *i*, *P*_*ji*_ is the probability of the connection from population *i* to population *j*, *W*_*ji*_ is the synaptic connection strength from population *i* to population *j* (see Methods for details). Of these, *d_*i*_* is estimated from the electron microscopy dataset, and *P*_*ji*_ and *W*_*ji*_ are estimated from the multi-patch synaptic physiology data. The model inputs consist of the constant baseline activity and the average activity of the L4 Exc neurons that are extracted from the Neuropixels data. The model’s output, in turn, is compared with the activity of each population in L2/3 derived from the same Neuropixels data. The weighted sum of squared error over the time course of the activity is the basis of the cost function (fit cost, Fig. 2).

It is well known that biological neural circuits may exhibit similar dynamical patterns with substantially different sets of parameters (Prinz et al., 2004). The key advantage of the SSN population model is its low computational cost, thus allowing one to explore multiple possible solutions and their statistical variance, rather than restricting the study to a single or a few possibly idiosyncratic solutions. By optimizing the model using a gradient-based minimization algorithm (see Methods for details) starting from 100,000 different initial parameter values for both Familiar and Novel sessions, we evaluated which solutions were suitable for subsequent analysis. We decided whether to accept the solutions based on a parameter cost cut-off (three standard deviations from the mean values obtained from the experimental data) and a fit cost threshold. Fig. 3A-D illustrates the distribution of the fit cost for each population, with an increasing cost to the right indicating a deteriorating fit. We investigated how solutions deviated from the target as a function of the fit cost and found that at the fit cost of 0.2 or above, solutions that fail to exhibit responses to image presentations start to appear (gray shading in Fig. 3A, C). The VIP population in the Familiar session is an exception; solutions with little visual responses are observed even at fit costs below 0.2. This is due to the low visual responses in the original target data (Fig. 3B bottom). Based on these observations, we decided to set our fit cost threshold to 0.2 while making sure that the solutions near the cut-off point still reasonably reproduce the target data traces (Fig. 3B, D).

**Figure 3:**
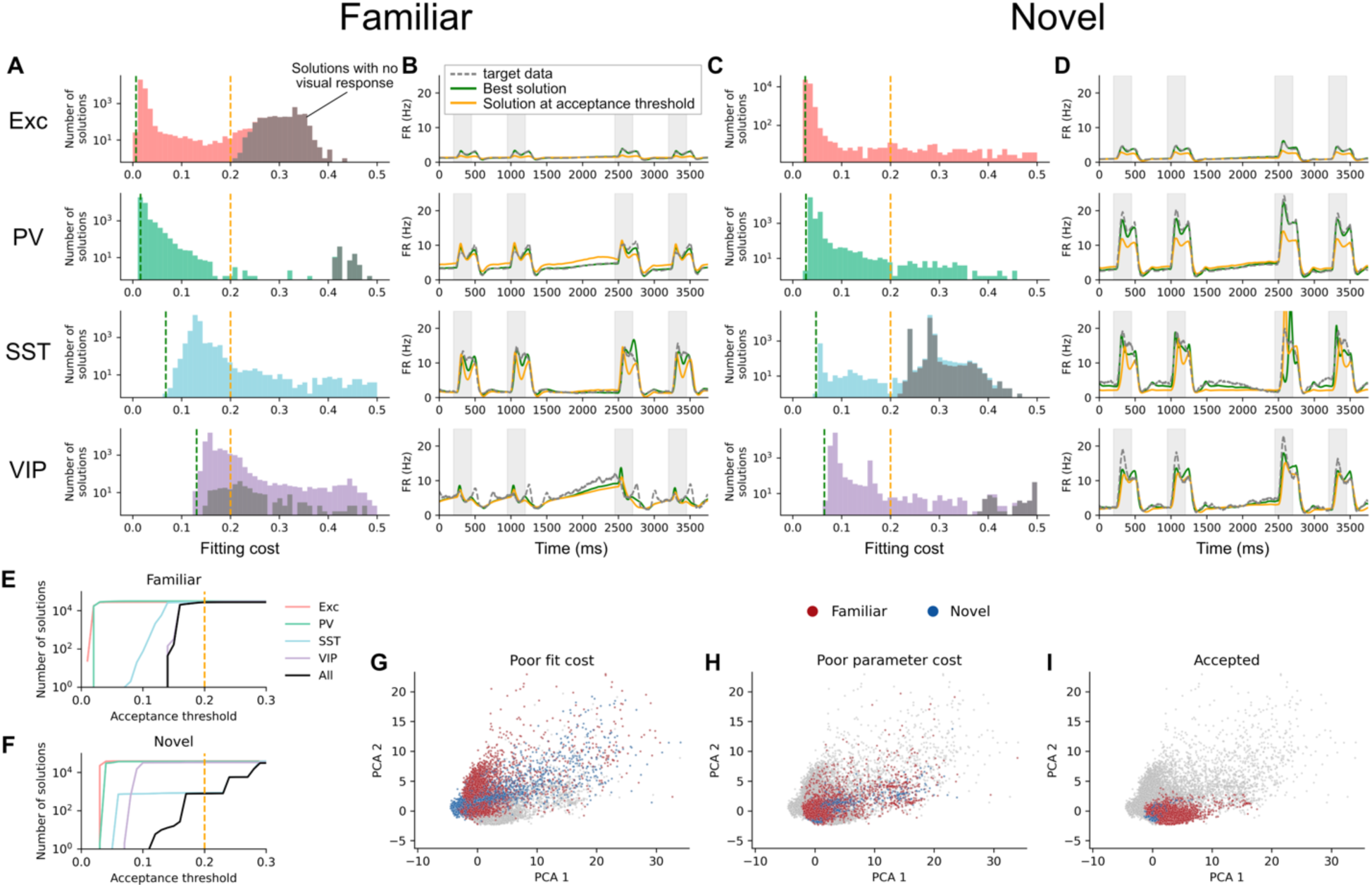
Statistics of the solutions and acceptance criteria. A: Histograms of the fitting cost values for all solutions passing the parameter cutoff criteria for the Familiar condition. The cost function values for the best solution and the solution near the acceptance threshold are indicated by green and orange dashed lines, respectively. The shaded area represents the number of solutions with no visual response (see Methods for details). B: Firing rate traces of the target data, the best solution, and a solution near the acceptance threshold for each population. C, D: Same as A and B, but for the Novel condition. E, F: Number of accepted solutions as a function of varying acceptance thresholds for the Familiar and Novel conditions, respectively. The black line indicates the number of accepted solutions wherein all populations passed the acceptance threshold. G-I: PCA projection of the parameters. Each panel highlights different categories of solutions with color: those with poor fit (G), those with poor parameter cost (H), and those that are accepted (I). Solutions with a fit cost greater than 0.3 for any population are excluded from the PCA analysis to enhance the visibility of the structure of the accepted solution. The accepted Familiar and Novel solutions form relatively tight clusters in the PCA space, providing a basis for subsequent LDA analysis.

Figures 3E and F show the number of acceptable solutions as a function of the acceptance threshold. The overall number of suitable solutions (i.e., solutions with all the populations accepted, indicated by the black line) remains relatively stable near the threshold, indicating that solution selection is not highly sensitive to a specific value of the threshold chosen.

To illustrate the selection of solutions, we performed the Principal Components Analysis (PCA) in the parameter space (Fig. 3G-I; the solutions with poor parameter cost, poor fit cost, and accepted solutions are highlighted). In total, we obtained 27,402 and 795 accepted solutions for the Familiar and Novel conditions, respectively. Solutions with poor parameter cost and poor fits have relatively large spread in the PCA space (Fig. 3G, H), but accepted solutions (Fig. 3I) are confined in a tight area in the PCA space, forming a relatively compact groups of points for the Familiar and for the Novel sessions. This again highlights the fact that a wide range of irrelevant parameters (Fig. 3H) can reproduce the population dynamics if the parameters are not constrained by the experimental data. The relatively simple structure of the accepted solutions in the PCA space sets a foundation for subsequent examination using Linear Discriminant Analysis (LDA).

### Coordinated changes in network parameters determine distinct dynamics in Familiar and Novel conditions

Having obtained tens of thousands of solutions that satisfactorily describe the distinct neural dynamics for the Familiar and Novel conditions, while using parameters consistent with the synaptic physiology and electron microscopy data, we set out to characterize the differences in these structural parameters that determine the differences in dynamics. One possible outcome is that certain parameters, such as synaptic weights of specific connection types, systematically change their values between the Familiar and Novel conditions. For example, certain connections could be stronger in the Familiar and weaker in the Novel condition, and vice versa for other types. However, we find this not to be the case. Fig. 4A shows histograms of all the parameters in the model for Familiar and Novel conditions. Note that we have included the scaling factor (*s* in Equation 1; see Methods for details) into the individual connection weights to do this analysis. Many parameters show heavily overlapping distributions, indicative of a wide range of possibilities that explains the transition with coordinated shifts in multiple parameters.

**Figure 4:**
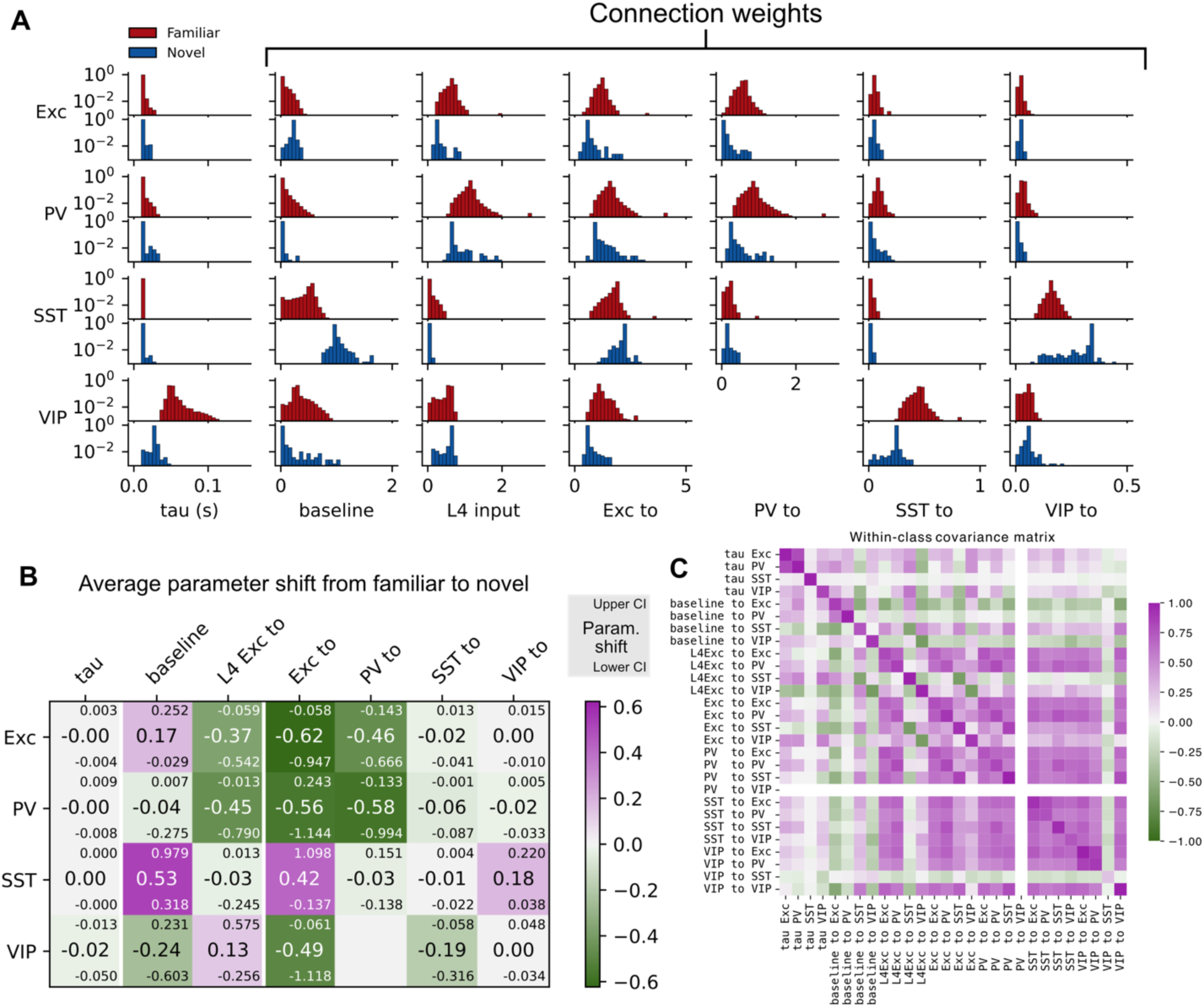
Summary of the parameter changes between the Familiar and Novel conditions. A: Histograms of model parameters for each condition. Although some parameter distributions show significant shift in their average values, most distributions overlap between the two conditions. B: Heatmap illustrating the average parameter shifts from Familiar to Novel condition. The confidence interval was calculated from the shifts in 10^6^ randomly sampled solution pairs. C: Covariance matrix depicting the relationships among the parameters within conditions.

To quantify these shifts, we calculated the mean difference in parameters between the Familiar and Novel conditions. Fig. 4B displays these shifts as a heatmap, with the 95% confidence intervals derived from 10^6^ randomly sampled solution pairs. Most connection parameters are weaker in the Novel condition compared to the Familiar one, particularly within the Exc and PV subnetworks. Note that this does not necessarily indicate an active weakening process; rather, it could reflect that strengthened connections are formed during training with Familiar images but are not utilized when processing Novel stimuli. On the other hand, the SST population receives stronger baseline, Exc, and VIP inputs in the Novel condition, marking an exception. While VIP to SST connection is stronger, SST to VIP connection is weaker in the Novel condition, hinting at a shift in the balance of interactions between these two populations.

Although these observations are instructive, it is critical to acknowledge that parameter changes are often correlated, adding another layer of complexity. Fig. 4C presents a within-class correlation matrix of the parameters, revealing substantial correlations among various connection weights. Our analysis benefits from a secondary measure that accounts for these complex correlation structures.

To capture the interdependencies between the parameters, we employed linear discriminant analysis (LDA) (Hastie et al., 2009). This technique identifies a hyperplane that maximizes the between-class variance while minimizing the within-class variance, thus optimally discriminating between the Familiar and Novel conditions (Fig. 5A; see Methods). The LDA-derived hyperplane exhibited a 100% separation accuracy between the conditions (Fig. 5B). However, visualizing the discriminative power across a high-dimensional parameter space (n=28) is non-trivial. To illustrate, Fig. 5C shows the projection of the data onto two representative axes, emphasizing how the correlation structure aids effective discrimination.

**Figure 5:**
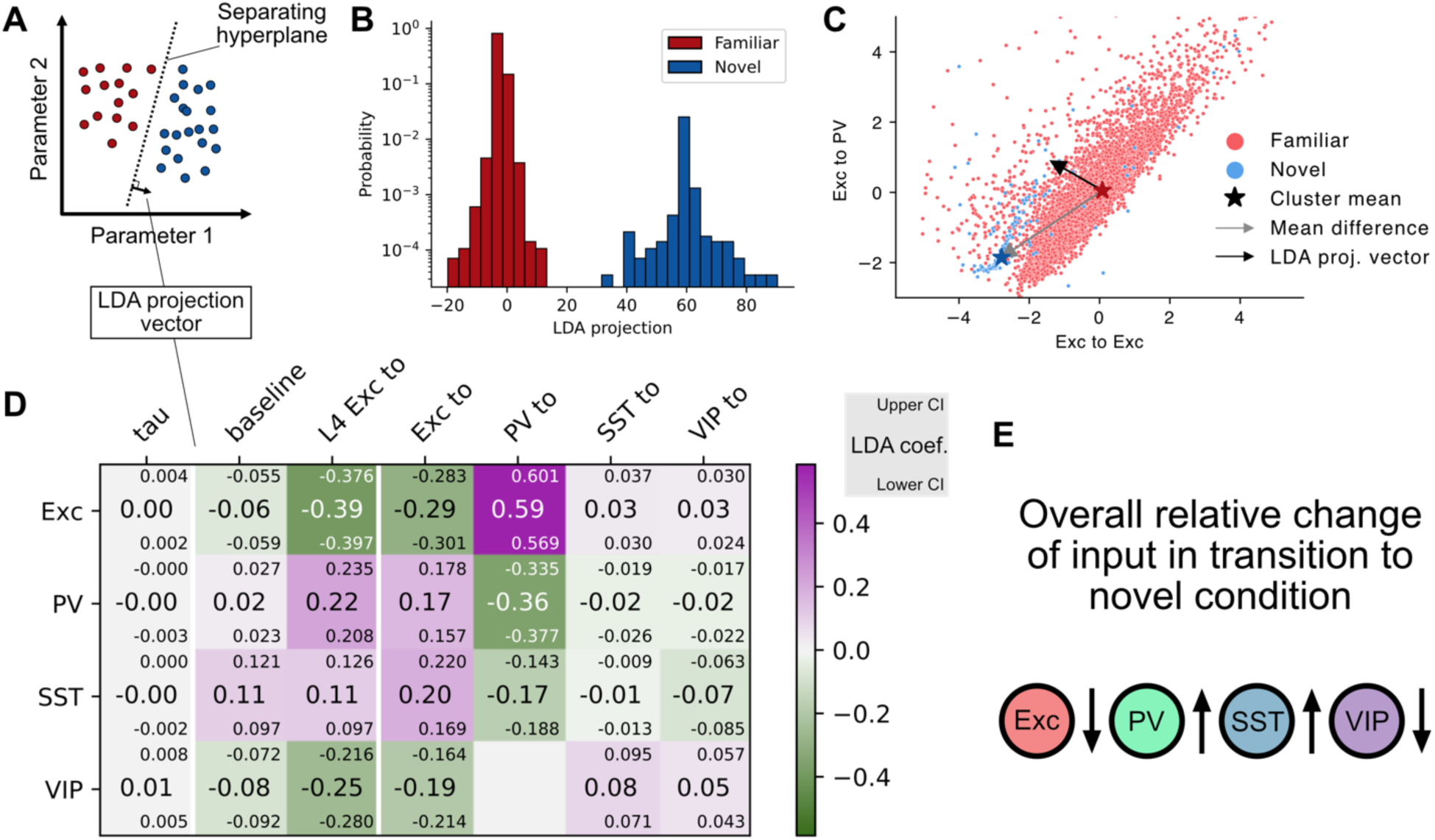
Statistical characterization of the accepted solutions using linear discriminant analysis (LDA). A: Schematic of the LDA procedure. It identifies a separating hyperplane in the parameter space, given the cluster labels. The vector orthogonal to the hyperplane is termed the LDA projection vector. B: Probability distribution of the number of solutions projected along the LDA projection vector. The familiar and novel conditions are completely separable. C: A visualization of the data points in the z-scored parameter space along two specific axes. Correlations between the parameters can be seen for both conditions and using such information enhances the separation effectiveness. D: LDA projection vector between the model solutions for the Familiar and Novel conditions, with colors indicating the weight of each parameter. A positive sign indicates that a higher parameter value is more attributable to the Novel condition, and vice versa. Meanwhile, a larger magnitude denotes a more substantial role in differentiating the conditions. E: Summary of relative changes in input to each neuronal population when transitioning from Familiar to Novel conditions. The transition is characterized by generally inhibitory input into the Exc and VIP populations and a generally excitatory input into the PV and SST populations.

An interesting observation arises when comparing the LDA vector to the mean shifts of individual parameters. For instance, although the average ‘Exc to PV’ parameter value decreases in the transition from Familiar to Novel conditions (Fig. 4B, gray arrow), the corresponding component in the LDA projection vector is positive (Fig. 5D, black arrow). This apparent contradiction is reconciled by considering the parameter’s correlation with other variables, such as ‘Exc to Exc.’ When networks share similar ‘Exc to Exc’ values, the ‘Exc to PV’ values are generally higher in the Novel condition. The LDA approach accurately captures this nuanced relationship, assigning a positive value to the ‘Exc to PV’ component.

As demonstrated above, the LDA projection vector offers a nuanced perspective on how parameters change between conditions, taking into account the correlation structures between them (Fig. 5D). Evaluation of the direction and magnitude of the LDA projection vectors yields several interesting results. First, the time constants of the neural populations contribute minimally to discrimination between the Familiar and Novel conditions. Second, outputs from Exc and PV populations as well as from the L4 Exc population (which plays the role of an external input into our model) all carry large coefficients, signifying their substantial role in transitioning from Familiar to Novel patterns of dynamics. Third, for the inputs into each population, all the excitatory connections exhibit the same sign, while all inhibitory connections show the opposite sign. This pattern indicates that the net change in the excitation-inhibition balance is more important for discriminating between the two conditions than identifying specific sources of excitation or inhibition.

While LDA effectively distinguishes between Familiar and Novel conditions, its output should be interpreted cautiously as an indicator of mechanistic changes in neural connectivity. First, parameter shifts in the LDA framework should be understood in relative terms, both within individual populations and across multiple populations. For instance, if excitatory inputs into the Exc population have an opposite sign to inhibitory inputs, this doesn’t necessarily mean there’s an absolute decrease in excitation and an increase in inhibition. Rather, it could indicate that both excitatory and inhibitory inputs have increased, but inhibition has done so more than excitation. Similarly, ‘negative input’ in the Exc population should be understood relative to other populations. In this context, a ‘negative input’ to Exc simply means its net change in inputs is less excitatory than what PV receives, irrespective of the absolute values. Indeed, the higher firing rates of the Exc population in the Novel condition (Fig. 1C) suggest that input into Exc population may actually be positively shifted. In this case, what the LDA vector suggests is that the PV and SST populations experience a larger positive shift of the inputs than the Exc population. Second, in the real biological system, the transition from ‘Familiar’ to ‘Novel’ may be driven by a limited set of solutions, not necessarily the full range that we observed. Consequently, LDA’s projection vector should be seen as a statistical descriptor of the model’s differential response to conditions, rather than a deterministic explanation for the changes of individual parameters.

Despite these limitations, an LDA analysis reveals a distinct pattern: The key factor for differentiating the networks is the relative net imbalance of the input. Specifically, upon transition to the Novel condition, Exc and VIP receive less excitatory input than PV and SST populations (Fig. 5E). These coordinated, relative shifts in connections are crucial in characterizing the disparities between the Familiar and Novel conditions.

### Connections to the VIP population have a unique contribution in separating image responses from non-image responses

The two major types of neural activity observed in the neural recordings we are using, either in Familiar or Novel conditions, are responses to images and responses when images are not shown (including image omissions). Are changes between the Familiar and Novel dynamics for these two response types determined by changes in distinct groups of the circuit parameters? To investigate this, we created additional target data sets of neural activity that combined the image and non-image response segments in different ways (Fig. 6A-D), resulting in two scenarios: 1) Familiar non-image responses (again, including omission responses) combined with Novel image responses, and 2) Familiar image responses paired with Novel non-image responses.

**Figure 6:**
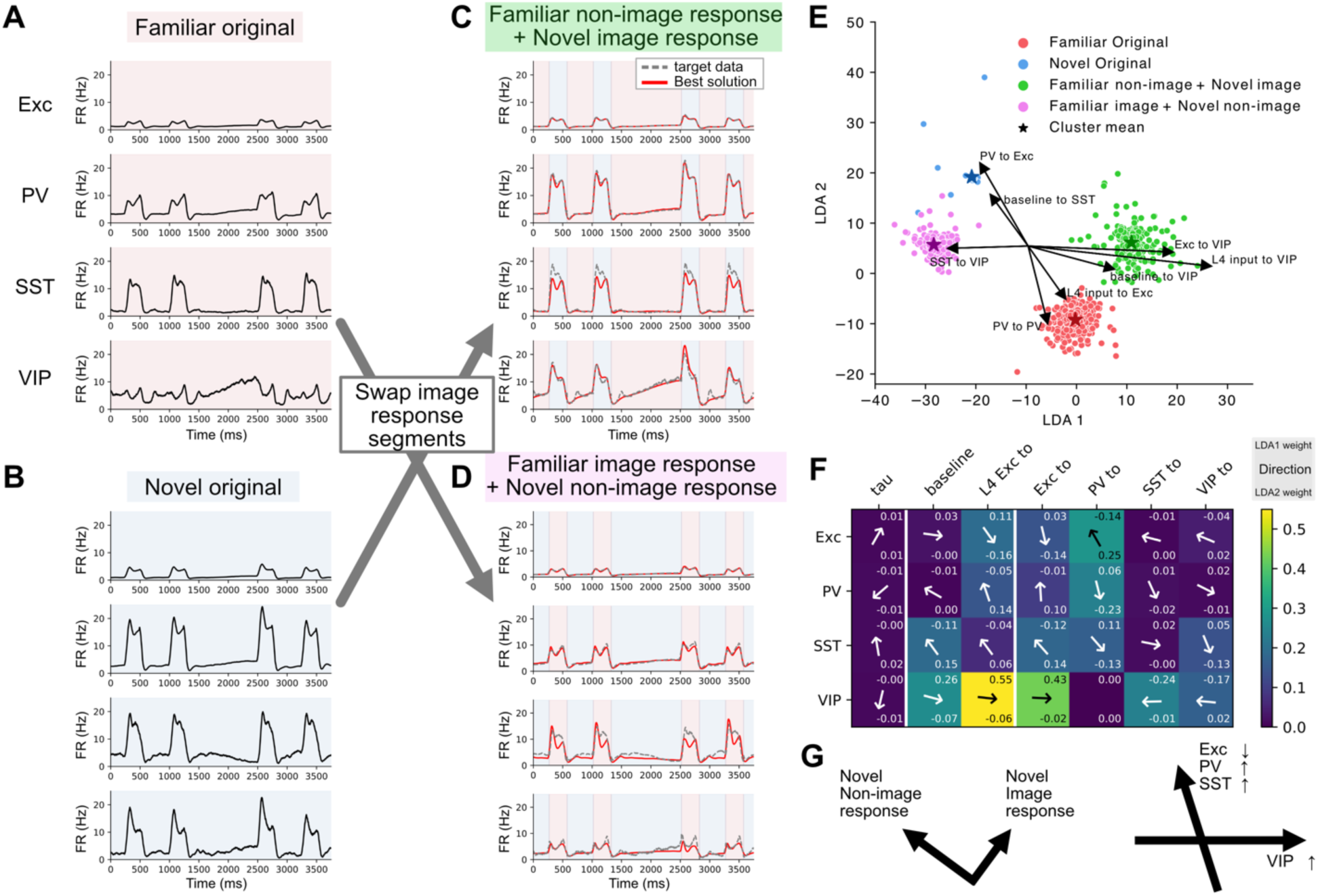
Target data manipulation to understand how different parts of the data affect the parameters. A, B: Firing rate traces of the Familiar and Novel datasets. C, D: Modified versions of the data wherein the image response and non-image response segments are swapped between sessions. E: Projection of the parameters of accepted solutions in the 2-dimensional LDA space, along with a biplot of the eight most influential parameters. F: Projection of each parameter into the 2-dimensional LDA space. The arrow denotes the direction, and the color indicates the weights of each element. G: Summary schematics of the target manipulation study. Each swapped session can be viewed as a change in certain segments from the original familiar session. When different segments are swapped, they form orthogonal vectors in the 2D LDA space (left). The summation of these changes corresponds to the total transformation of the firing rate traces. Notably, alterations to inputs into the VIP population uniquely result in horizontal movement in the LDA space, corresponding to the swapping of different segments (right).

Applying the same procedure as in the previous sections to these new data sets (together with the original all-Familiar and all-Novel target datasets) and performing multidimensional LDA, we found that the parameter sets formed four distinct clusters in the LDA space – one cluster for each of the target datasets – resembling a parallelogram (Fig. 6E). This geometric arrangement suggests that the changes in firing rate traces can be broken down into different components, which, when combined, yield the overall transition effect. Importantly, the projection of several of the top contributing parameters on the LDA space (Fig. 6E) highlights inputs into the VIP population (horizontal axis in the map) as a critical factor in characterizing the difference between the image response part and the non-image response part.

Fig. 6F shows the projection of each parameter onto the 2D LDA space in Fig. 6E, illustrating each parameter’s contribution to the change in the LDA space. We find that any substantial inputs into the Exc, PV, and SST populations lie along the axis that connects the original all-Familiar and all-Novel networks, while the inputs into the VIP population lie along the horizontal axis (summarized in Fig 6G). This confirms the observation made in Fig. 6E, using the top 8 contributing parameters, but in a more concise manner. The direction of the change of the VIP population implies a necessity for high levels of input into the VIP population to achieve the Familiar non-image response and Novel image response.

### Simulated Novel network has higher gain and earlier saturation compared to Familiar network

To understand how our SSN models respond to input stimuli that they were not optimized for, we tested the network response with artificial square-pulse stimuli with various amplitudes. The amplitude of the stimuli was varied while the baseline firing rates were kept constant (Fig. 7A). Fig. 7B, C shows responses of example solutions for the Familiar and Novel cases to inputs with various amplitudes. In general, the response amplitude increases as a function of the stimulus amplitude, but the temporal structure and relative proportion of the responses of each population are different between these two networks. Fig. 7D, E characterizes the responses of each neural population to a range of input amplitudes for all accepted Familiar and Novel solutions.

**Figure 7:**
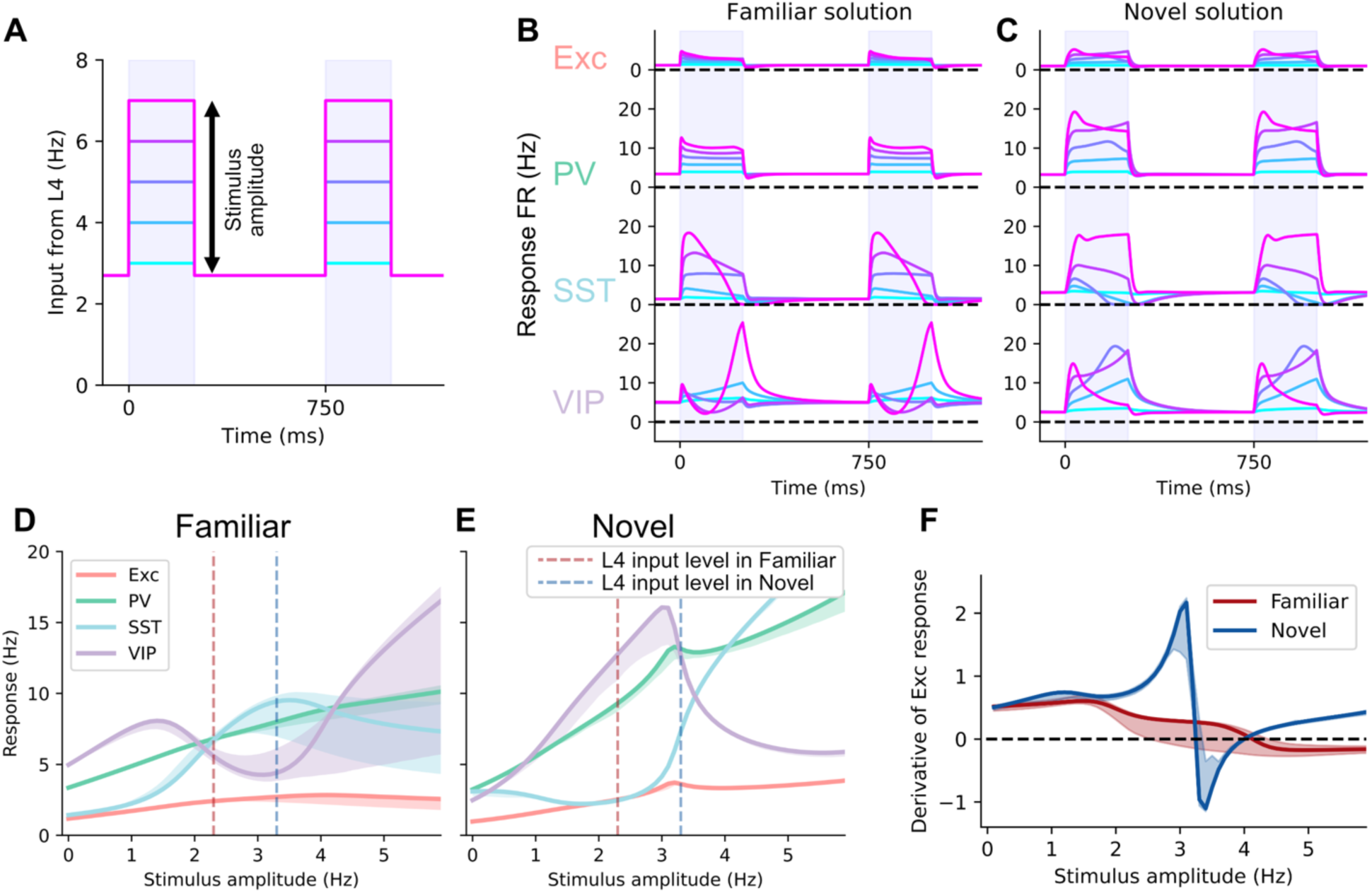
The networks in the Novel condition have higher gain and earlier saturation compared to those in the Familiar condition. A: Illustration of the stimuli applied with the baseline firing rate fixed at 2.7 Hz for both sessions and the stimulus amplitude indicating deviation from 2.7 Hz. B, C: Population responses to the stimuli with different amplitudes for example solutions of the Familiar and Novel conditions, respectively. The colors of the traces correspond to those shown in (A). D, E: Population responses as a function of the stimulus amplitude for all the solutions in the Familiar and Novel conditions, respectively. The dashed lines indicate the stimulus levels of the experimental Familiar and Novel sessions (activity of the L4 Exc population). F: The derivative of the Exc population responses as a function of the stimulus amplitude (stimulus gain) for the Familiar and Novel solutions.

A particularly interesting observation was made when analyzing the amplitude response of each population. Regardless of whether the network was coming from the Familiar or Novel pool of solutions, there was a region with weaker stimuli where the response of the VIP population exceeded that of the SST population. However, as the stimulus strength increased, the response of the SST population surpassed that of the VIP population (Fig. 7D, E). This finding aligns with the results of a previous report, which shows that L2/3 VIP neurons are tuned to low-contrast stimuli, and SST neurons are tuned to high-contrast stimuli (Millman et al., 2020). Importantly, the extent to which VIP activity dominates over SST is greater in the Novel condition, corroborating our observations in Fig. 4B regarding the altered balance between these populations.

The Familiar networks displayed smooth, consistent responses to changes in the input firing rates. Conversely, the Novel networks exhibited an abrupt change or ‘kink’ in the responses at an input amplitude of around 3 Hz. And beyond this point, the firing rates of the Exc population slightly decrease as the stimulus amplitude increases up to ∼4 Hz. This breaks the 1-to-1 relation between the input stimulus amplitude and the output firing rates of the Exc population and thus makes it infeasible to estimate the input amplitude from the output amplitude.

It is important to note that the stimulus amplitudes (i.e., the firing rates of the L4 Exc populations, providing the major input to L2/3) recorded during the experiments were different between the Familiar and Novel conditions: 2.3 Hz for the former vs. 3.3 Hz for the latter. The amplitude of the stimulus could be weaker with reduced visual stimulus contrast. However, because we used full-contrast stimuli in the experiment, it is not possible to further increase the L4 activity by increasing the visual stimulus contrast. Therefore, we consider amplitudes significantly larger than these measured values (e.g., > 4 Hz; note that this is the mean firing rate for the whole population) to be physiologically unrealistic.

Lastly, Fig. 7F plots the derivative of the firing rate of the Exc population as a function of the stimulus amplitude, which effectively represents the gain (input/output ratio) of the network. Here we observe that while the Novel networks showed a slightly higher gain overall, there was an anomaly around the input amplitude of ∼3 Hz, which corresponds to the aforementioned kink in the output of the neural populations. This implies that the Novel network, while having a higher gain to the stimulus amplitude, also exhibits an earlier saturation of firing rates.

## Discussion

We investigated the changes in the cortical network when an animal is exposed to novel stimuli, using an SSN model. Our approach involved extensive sampling of the parameters that are constrained by various integrated data sources. Our analysis revealed a general trend of weakening connectivity when transitioning from the Familiar to Novel conditions. This weakening was not uniform, and it was accompanied by a notable shift in the balance between SST and VIP populations. LDA analysis added granularity by effectively discriminating the two conditions based on the relative net input changes across populations. Specifically, in the Novel condition, PV and SST populations experienced overall more excitatory influences compared to Exc and VIP populations. We also found that the input into the VIP population plays a unique role in separating image response and non-image response. Lastly, using stimulation at varying amplitudes, we observed that the network in the Novel condition exhibited higher gain and earlier saturation to the stimulus amplitude compared to the network in the Familiar condition.

### Data integration with the SSN model yields networks that generate observed dynamics with realistic parameters

We sought to leverage the strength of the SSN model by integrating it with various publicly available data. The Neuropixels Visual Behavior dataset, used for both input and target data, includes recordings from many neurons across many mice engaged in a systematically designed active behavior. In addition, this dataset provides information about the activity of subclasses of inhibitory neurons (PV, SST, and VIP), identified with waveform clustering and opto-tagging. The fine temporal resolution of the Neuropixels electrophysiology resolves spikes of the recorded neurons, allowing us to operate quantitatively with the directly measured firing rates, rather than secondary measures of neuronal output like the calcium signal. Most of the model parameters, primarily connectivity parameters, were systematically constrained by the other available datasets. The net influence between populations was estimated based on connection probability, connection strength, and the abundance of the neurons in each subclass. Multi-patch synaptic physiology data (Campagnola et al., 2022; Seeman et al., 2018) and the MICrONS electron microscopy dataset (The MICrONS Consortium et al., 2021) provided these values at the granularity of subclass level in each layer of V1—a level of detail that was unavailable prior to their publication. We applied the constraints based on these data to the connection parameters while allowing for changes that could reasonably be expected, given the measured variability in the data.

### Extensive sampling of the solutions finds coordinated shift of the model parameters between the Familiar and the Novel conditions

We concentrated on obtaining extensive samples of the possible set of parameters instead of analyzing a small number of solutions. Even with the constraints on the network parameters determined by the biological data, having redundant solutions with variable parameter sets is unavoidable. Thus, we employed a systematic optimization strategy that allowed us to characterize the solution space capable of producing activity patterns that quantitatively match the experimental data.

As summarized in Figure 8, our results indicate a coordinated shift in connection strengths between the Familiar and Novel conditions. We observed two major shifts: a general weakening of synaptic connections and a reconfiguration of inhibitory control, pivoting from SST to VIP populations. This latter shift aligns with the expanded VIP-dominant region observed in the gain curve (Fig. 7E) during Novel conditions.

**Figure 8:**
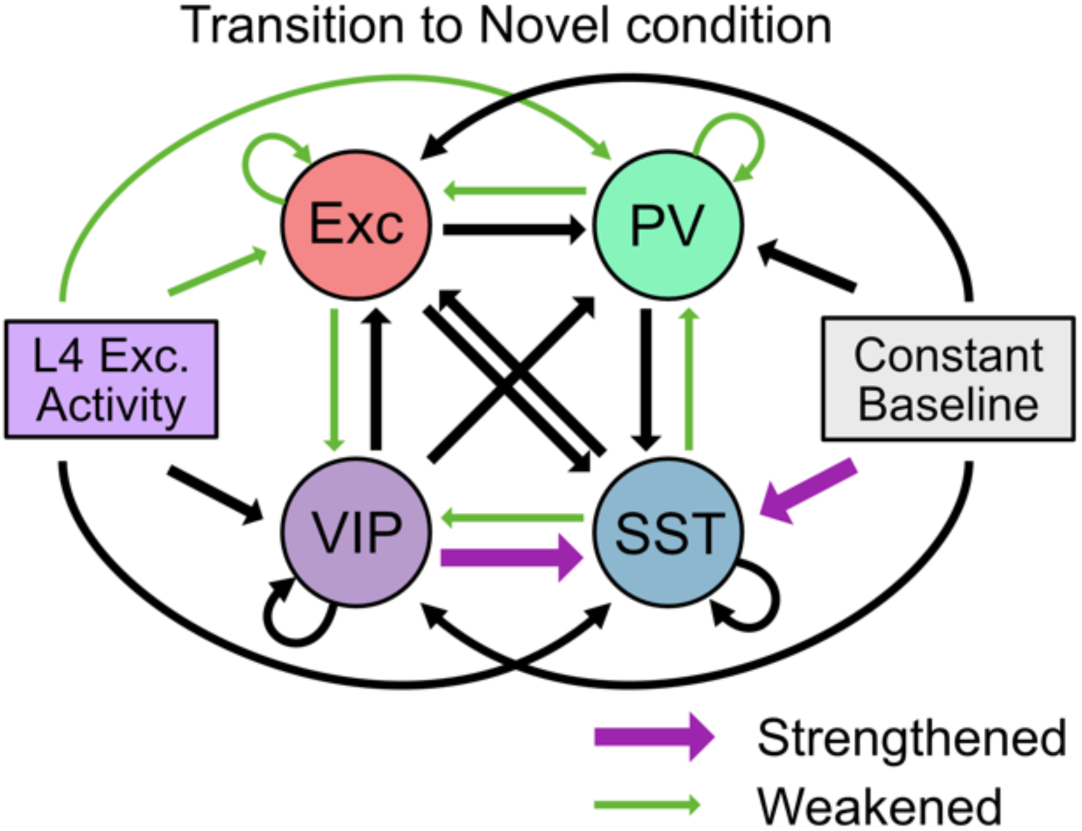
Summary schematic of the transition from the Familiar to the Novel network. This figure illustrates key changes in the connections in our model networks. Connections exhibiting changes with wholly positive confidence intervals are labeled as ‘strengthened,’ while those with wholly negative confidence intervals are labeled as ‘weakened’ and further emphasized by colored arrows (see Fig. 4B). The transition from the Familiar to the Novel condition is characterized by general weakening of the connections, with increased control of the SST population by the VIP population.

Several mechanisms could potentially explain the observed changes in the connection strength. The general weakening of the connections could be attributed to the utilization of a naïve neural pathway that is not facilitated when processing Novel, untrained stimuli, compared to more familiar ones. Other factors such as synaptic plasticity across various time scales and neuromodulatory influences could also be contributing to changes in connection strengths, not limited to the observed weakening, representing additional areas for future investigation.

Our findings thus raise additional questions about the complex neural dynamics in V1, offering a basis for comparison with other modeling studies. For example, our approach contrasts with a recently published study by Aitken et al. (2023), which also employs the same experimental paradigm developed by Garrett et al. (2020, 2023). Aitken et al. focus on elucidating a biologically plausible mechanism of synaptic plasticity, capable of reproducing novelty effects across different timescales, while also simulating individual neurons to describe their responses and diversity; however, they do not extensively seek the specific circuitry changes or their parametric details. On the other hand, our study aims to identify potential modifications in the structure of the V1 circuit in a population model, without making assumptions about the specific mechanisms responsible for these changes. These distinct but related objectives demonstrate that our respective works offer complementary insights into V1 neural dynamics.

### Unique role of inputs into VIP populations in separating image and non-image responses

The role of inputs into VIP populations appears to be unique, as they contribute to the separation of network parameters tuned to synthesized target data with swapped image and non-image responses. This indicates that these inputs may be more dependent on the specific timing of the stimuli, implying they receive more nuanced inputs that can vary within a condition. However, given that our study does not include time-varying external inputs beyond layer 4, we are unable to fully describe how external influences from the rest of the brain impact the network dynamics of the modeled area.

Further investigations, potentially utilizing more elaborate models, will be necessary to explore these contributions because novelty signal is known to affect many interconnected brain areas (Knight, 1996). Potential pathways for such top-down modulation could include inputs from higher-order visual areas (Glickfeld & Olsen, 2017) and the cingulate cortex (Zhang et al., 2014). Studies in other brain areas identify the involvement of neuromodulators such as acetylcholine, noradrenaline, and dopamine (Kafkas & Montaldi, 2018; Ranganath & Rainer, 2003). While the changes in the circuit parameters uncovered by our modeling study here may very well be due to the action of such neuromodulators on the neuronal and synaptic properties, our model is agnostic to the specific mechanisms of these changes. Future studies should consider including these neuromodulatory effects by investigating such mechanisms experimentally and including them explicitly in models.

### Trade-off between higher gain and early saturation in the Novel network

Our findings suggest that the novel network might have a higher gain of firing rates potentially at the cost of early saturation to the input amplitude. In the familiar network, the gain was lower but exhibited a steady increase in the firing rate over the physiological range (up to ∼4 Hz of stimulus amplitude). Conversely, the novel network exhibited a ‘kink’ in the firing rate curve, beyond which the gain becomes negative even within the physiological range of stimulus amplitude. This property may enhance sensitivity to low-amplitude stimuli, but impair the distinction of input amplitude beyond the ‘kink’. While our model provides an insightful observation, it is important to recognize that this predicted characteristic remains to be validated in an experiment. One potential approach would be to present stimuli of varying contrasts and examine whether the network’s contrast sensitivity changes in the Novel condition.

## Conclusion

Overall, our study highlights the potential of integrating diverse data modalities into a unifying modeling framework, which gave insights into how the network generates complex dynamics observed in experiments. In addition to the coordinated shift of the parameters between the Familiar and Novel conditions, our work underscores the unique role of the VIP population in encoding different inputs (i.e. image vs. non-image) and proposes potential trade-offs in neural processing under varying contexts. Future work should investigate aspects unattainable with the current modeling framework, such as the inclusion of external stimuli beyond the layer 4 population, or the simulation of individual spiking neurons to capture heterogeneity within a population, for example, using large-scale networks of spiking point or multi-compartmental neuron models and similarly integrating multi-modal experimental data constraints (Billeh et al., 2020; Haufler et al., 2023).

## Methods

### Target and input firing rate traces for the model

We extracted the firing rate traces of populations of neurons from the Allen Institute Visual Behavior Neuropixels data (https://portal.brain-map.org/explore/circuits/visual-behavior-neuropixels), which served as both the target data and the input for our modeling. Neurons are segregated into four putative populations: Exc, PV, SST, and VIP populations. The SST and VIP populations were identified by opto-tagging (using the method described in https://allensdk.readthedocs.io/en/latest/_static/examples/nb/ecephys_optotagging.html). The neurons that were not opto-tagged were classified using their waveforms. If the waveform duration is smaller or larger than 0.4 ms, they are considered putative PV (fast-spiking inhibitory) or Exc neurons, respectively.

The neurons were separated into layers based on the cortical depth of the recording. The cortical depth was estimated by the vertical distance from the recorded neuron closest to the surface. Neurons with depth less than 200 µm were identified as layer 2/3 neurons and neurons with depth more than 200 µm and less than 300 µm were identified as layer 4 neurons. The Exc, PV, and SST neurons were taken only from L2/3 of V1, but the VIP neurons in the Novel condition were taken from all layers of V1 because of the small number of opto-tagged neurons in layer 2/3. Table 1 summarizes the number of identified neurons for each population and the number of mice from which the data were collected. We first calculated the average population activity for each mouse and then averaged across the mice. We used the mouse-wise standard deviation as the error for fitting.

**Table 1:** Number of identified neurons and the number of mice the neurons are recorded from. (*Note that there were only 3 VIP neurons recorded in layer 2/3 in the Novel condition, so we added neurons that were recorded in other layers of V1.)

| Cell subclass | Number of identified neurons (Familiar) | Number of mice (Familiar) | Number of identified neurons (Novel) | Number of mice (Novel) |
| --- | --- | --- | --- | --- |
| Exc | 758 | 34 | 519 | 35 |
| PV | 167 | 33 | 120 | 34 |
| SST | 12 | 8 | 10 | 5 |
| VIP | 6 | 3 | 5 * | 4 |
| L4 Exc | 595 | 34 | 454 | 35 |

The input stimulus to the model is taken from the same Neuropixels dataset. These neurons were chosen from those that were identified as the putative Exc neurons but from layer 4 (L4 Exc).

### Four-population Stabilized Supralinear Network (SSN) model

We used the stabilized supralinear network (SSN) model (Rubin et al., 2015) to capture network dynamics. The model consists of four interacting populations (Exc, PV, SST, and VIP of V1 layer 2/3 of the mouse), receiving constant baseline inputs and inputs from L4 Exc population. The model equations are:

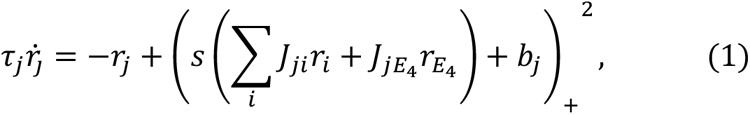

where *r*_*j*_ is the firing rate of population *j* (*j* ∈ *Exc*, *PV*, *SST*, *VIP*), *τ*_*j*_ is a decay constant of the population *j*, *s* is a scaling factor of the connectivity, *J*_*ji*_ is the strength of the connection from population *i* to population *j* (*i* ∈ *Exc*, *PV*, *SST*, *VIP*, *L*4 *Exc*), and *b*_*j*_ is a baseline input into population *j*. The *x*_+_ indicates rectification of *x* at zero: *x*_+_ = *x* if *x* > 0; = 0 otherwise.

The connection between populations is estimated using 1) the probability of connections between two populations (Fig S1 of (Campagnola et al., 2022)) in L2/3; the average connectivity weights between two populations ((Campagnola et al., 2022), https://github.com/AllenInstitute/aisynphys/blob/figure-notebooks/doc/figure_notebooks/figure_03_strength_kinetics.ipynb; From section D) Resting state amplitude of postsynaptic responses among cell classes); and the abundance of each population estimated by the MICrONS EM data (https://www.microns-explorer.org/cortical-mm3). The relative weights between the population were estimated with the following relationship:

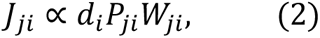

where *J*_*ji*_ is the connection strength from population *i* to population *j* used in the SSN model, *d*is the density of the neuron, *P*_*ji*_ is the connection probability between neurons in given populations, and *W*_*ji*_ is the average synaptic strength between connected neurons in given populations. Table 2 summarizes the estimated values. Supplementary Data 1 lists all the source information and outlines how the matrix elements are calculated. The overall scale of these values is not constrained, thus we considered it as a parameter (*s*) of the model. Because of this scaling, it creates redundancy of the model’s behavior with some parameter combinations. For example, when all *J*_*ji*_ are doubled and *s* is halved, the model’s behavior is unchanged. To avoid analyzing these redundant models separately, we incorporated the scaling factor into each connectivity matrix element.

**Table 2:** Data-based estimates of the connection strengths (*J*_*ji*_) and their standard errors.

| Target\Source | L4 Exc | Exc | PV | SST | VIP |
| --- | --- | --- | --- | --- | --- |
| Exc | $0.024 \pm 0.013$ | $0.029 \pm 0.013$ | $-0.010 \pm 0.004$ | $-0.0024 \pm 0.0009$ | $-0.00046 \pm 0.00029$ |
| PV | $0.27 \pm 0.13$ | $0.40 \pm 0.15$ | $-0.016 \pm 0.005$ | $-0.0020 \pm 0.0008$ | $-0.0005 \pm 0.0005$ |
| SST | $0.003 \pm 0.004$ | $0.19 \pm 0.08$ | $-0.0036 \pm 0.0022$ | $-0.0006 \pm 0.0003$ | $-0.007 \pm 0.003$ |
| VIP | $0.03 \pm 0.03$ | $0.28 \pm 0.11$ | N/A | $-0.0074 \pm 0.0025$ | $-0.0010 \pm 0.0009$ |

### Fitting procedures

To obtain models that achieve both good fits and small parameter deviation, we used a compound cost function that has two elements, as follows:

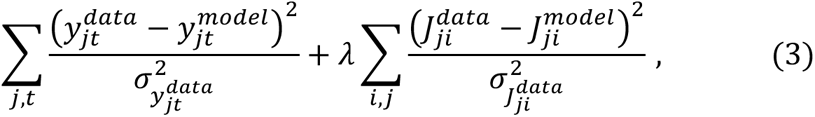

where 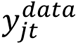 is the firing rate of population *j* at time *t* in the target data, 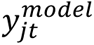 is that predicted by the model, 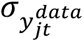 is the standard error of the firing rate of population *j* at time *t* in the target data, *λ* is a hyperparameter that sets the relative weight of the parameter cost, 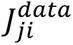 is an estimated influence from population *i* to population *j* based on experimental data, 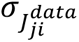 is the estimated uncertainty of it, 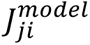 is the influence from population *i* to population *j* as a model parameter. The first term is the fit cost, and the second term is the parameter cost. We have explored the weight of the parameter cost *λ* and found that *λ* = 10 achieves a good balance between the quality of fit and the deviation of the parameters.

Many potential parameter combinations can produce reasonable results in this setup. To analyze the collection of such solutions, we decided to use 100,000 initial parameter sets and optimize each of them to obtain various solutions. The initial parameters were drawn from a uniform distribution in either the linear space or the log space (see the ‘Sampling method’ column in Table 3). Each set of the initialized parameters was optimized using the above cost function with the ‘migrad’ method (a gradient-based optimization algorithm) of the ‘iminuit’ package (https://iminuit.readthedocs.io; (James & Roos, 1975)) to optimize the function.

**Table 3:** Description of model parameters.

| Parameter | Description | Range | Number of parameters | Sampling method |
| --- | --- | --- | --- | --- |
| $\tau_j$ | Firing rate decay constant | 0.01-1 (seconds) | 4 | Log |
| $b_j$ | Baseline input into each population | 0-10 | 4 | Linear |
| $J_{ji}$ | Influence from $i$ to $j$ | 0.1-10, multiplied by the estimate from data (Table 2) | 19 (sources: L4 Exc, Exc, PV, SST, VIP; targets: Exc, PV, SST, VIP; according to the data, PV to VIP connection is not present) | Log |
| $s$ | Scaling factor of the influence matrix | 1-100 | 1 | Log |

Once the solutions were obtained, we filtered them for subsequent analyses. First, solutions with poor parameter costs were removed. The influence matrix parameters with values 3 standard errors above or below the data-based estimates were considered poor parameters, and the solutions that contain such a parameter were excluded. The histograms in Fig. 3A and C were plotted after excluding solutions with this parameter cost criterion. The threshold for the fit cost for each population was set to 0.2 as described in Results. We found that the solutions with a fit cost above this value start to have no visual responses. To determine whether a population has a visual response, we took the ratio of the firing rates between the image response period (70-305 ms after the stimulus onset; 70 ms is when the image response starts, and 305 ms is when the image response ends) and non-image response period (305-820 ms after the stimulus onset; 820 ms is when the response to next stimulus starts (the stimulus period is 750 ms)) and determined whether their ratio is more than 1.2.

### Target firing rate manipulation

To understand whether each parameter contributes differently to different temporal parts of the data, we decided to fit the model to synthesized data by combining different temporal parts of the existing data. We separated the firing rate traces of each population into the image response part (70-350 ms after the stimulus onset; 350 ms is when the firing rates go back to baseline firing rates from an undershoot) and the non-image response part (350-820 ms after the stimulus onset) and created synthesized data with 1) Familiar non-image response and Novel image response and 2) Familiar image response and Novel non-image response. After combining the data, a Gaussian temporal filter with a 20-ms width was applied to smooth the firing rates at the transition points. For this analysis, we used 10,000 initial parameter sets instead of 100,000 to reduce the computational cost.

## Code Availability

The code supporting the findings of this study is openly available on GitHub at the following repository: https://github.com/AllenInstitute/visual_behavior_ssn_modeling. The repository contains all the necessary code scripts and data processing modules used for the computational models and analyses described in this manuscript.

## Acknowledgments

We thank Kyle Aitken, Saskia de Vries, Kenta Hagihara, Christof Koch, and Stefan Mihalas for their feedback on this study. We thank the founder of the Allen Institute, Paul G. Allen, for his vision, encouragement, and support. This work was supported by the National Institute of Biomedical Imaging and Bioengineering of the National Institutes of Health under Award Number R01EB029813 and the National Institute of Neurological Disorders and Stroke of the National Institutes of Health under Award Numbers R01NS122742 and U24NS124001. The content is solely the responsibility of the authors and does not necessarily represent the official views of the National Institutes of Health.

## Notes

### Competing Interest Statement

The authors have declared no competing interest.

